# Predicting audiovisual speech: Early combined effects of sentential and visual constraints

**DOI:** 10.1101/360578

**Authors:** Heidi Solberg Økland, Ana Todorović, Claudia S. Lüttke, James M. McQueen, Floris P. de Lange

## Abstract

In language comprehension, a variety of contextual cues act in unison to render upcoming words more or less predictable. As a sentence unfolds, we use prior context (*sentential constraints*) to predict what the next words might be. Additionally, in a conversation, we can predict upcoming sounds through observing the mouth movements of a speaker (*visual constraints*). In electrophysiological studies, effects of visual salience have typically been observed early in language processing, while effects of sentential constraints have typically been observed later. We hypothesized that the visual and the sentential constraints might feed into the same predictive process such that effects of sentential constraints might also be detectable early in language processing through modulations of the early effects of visual salience. We presented participants with audiovisual speech while recording their brain activity with magnetoencephalography. Participants saw videos of a person saying sentences where the last word was either sententially constrained or not, and began with a salient or non-salient mouth movement. We found that sentential constraints indeed exerted an early (N1) influence on language processing. Sentential modulations of the N1 visual predictability effect were visible in brain areas associated with semantic processing, and were differently expressed in the two hemispheres. In the left hemisphere, visual and sentential constraints jointly suppressed the auditory evoked field, while the right hemisphere was sensitive to visual constraints only in the absence of strong sentential constraints. These results suggest that sentential and visual constraints can jointly influence even very early stages of audiovisual speech comprehension.

## 1. Introduction

If, during an English conversation, you see your friend put her upper teeth against her lower lip, you would know which kind of speech sound to expect next: a labiodental fricative consonant, i.e. either /f/ or /v/. These two different speech sounds share the same *viseme* (the facial gesture we can see when someone utters a speech sound). Visemes have been suggested to function as predictive cues to upcoming acoustic speech (Peelle & Sommers, 2015). However, sentence context may also provide you with valuable information about upcoming speech. For example, your friend might say, “*The firewood immediately caught […]*”. Such a context enables you to draw on both your linguistic and world knowledge (e.g., about the expression “to catch fire” and about the properties of firewood). In this case you might be able to predict “caught fire” as a likely ending. Sentential constraints could thus also serve as predictive cues (Altmann & Mirković, 2009). In the current study, we asked whether visual and sentential constraints jointly influence language comprehension and, if so, when they do so. Specifically, can contextual cues (e.g., sentential information about upcoming “fire”) modulate uptake of visual cues (e.g., mouth movements about the upcoming /f/-/v/ viseme) early in the comprehension process?

During speech recognition, visual information often precedes auditory information (Chandrasekaran, Trubanova, Stillittano, Caplier, & Ghazanfar, 2009). Visemes can communicate information about the place of articulation of a speech sound earlier and more efficiently than the acoustic signal alone (Jesse & Massaro, 2010). While visemes can therefore be used to predict upcoming acoustic speech, the degree to which they can do so depends on the level of certainty with which they signal (classes of) speech sounds. We will refer to this as *viseme salience*. The viseme for /f/ and /v/ is highly salient, as critical aspects of the articulation of these sounds are clearly visible. Visemes such as those belonging, for example, to velar consonants (/ɡ/ and /k/) are less salient, since the constriction that produces the sound is not visible. The available visual information associated with velar consonants is consistent with sounds produced at other posterior places of articulation. Viseme salience is reflected in an early neuronal response: the auditory N1 peaks earlier and has a lower amplitude with more salient visemes (Baart, 2016; Klucharev, Möttönen, & Sams, 2003; Wassenhove, Grant, & Poeppel, 2005), a phenomenon we will refer to as the *viseme effect*.

The auditory N1 is an event-related potential (or event-related field) peaking around 100 ms following a change in the acoustic environment (Hari et al., 1987). It is also considered to reflect early stages of language processing such as prelexical acoustic-phonetic analysis (Bien, Lagemann, Dobel, & Zwitserlood, 2009; Sjerps, Mitterer, & McQueen, 2011; Toscano, McMurray, Dennhardt, & Luck, 2010). Although traditionally seen as a stimulus-driven response to an auditory event, the N1 has also been found to be sensitive to higher-level cognitive states like attention and expectation (Hari et al., 1987; Näätänen & Picton, 1987; Todorovic, Ede, Maris, & Lange, 2011). A possible interpretation of the viseme effect, then, is that visemes serve to influence the probability of which sounds might be encountered next, with more salient visemes enabling more accurate predictions. Many viseme studies, however, used isolated phonemes or syllables (Arnal, Morillon, Kell, & Giraud, 2009; Arnal, Wyart, & Giraud, 2011; Baart, 2016; Wassenhove et al., 2005), thus trading off tighter experimental control for slightly lower ecological validity. One study that did employ an audiovisual paradigm with speakers uttering entire sentences, interestingly, found a reverse viseme effect (a stronger N1 for high visual saliency) confined to the right hemisphere (Brunellière, Sánchez-García, Ikumi, & Soto-Faraco, 2013).

In addition to viseme salience, there is considerable evidence to suggest that we are able to use grammatical, semantic and pragmatic sources of information in sentences to make predictions about what words will come next in those sentences (Altmann & Kamide, 1999; Arai & Keller, 2013; Dahan & Tanenhaus, 2004; Federmeier, 2007; J. L. Miller, Green, & Schermer, 1984; Pickering & Garrod, 2013; van Alphen & McQueen, 2001; Van Berkum, Brown, Zwitserlood, Kooijman, & Hagoort, 2005). In electrophysiological studies, sentence context predictions have been shown primarily to modulate the N400 (Kutas & Hillyard, 1980). The N400 is a negative deflection in the event related potential (ERP) that peaks approximately 400 milliseconds after word onset, but is often visible from 250ms onwards. The N400 to a word that is not predicted or is unexpected will have a higher amplitude than if the word was predicted or expected (Kutas & Federmeier, 2011; Lau, Phillips, & Poeppel, 2008).

Are the abilities to use visual and sentential information two sides of the same coin, that is, do they reflect the same predictive process? Before we attempt to answer this question, it is important to be clear what we mean by “prediction”. Different mechanisms that could support prediction have been proposed, including those based on pre-activation (DeLong, Urbach, & Kutas, 2005), on Bayesian principles (Kuperberg & Jaeger, 2016; Norris, McQueen, & Cutler, 2016), on generative models (Pickering & Garrod, 2007, 2013), and on predictive coding (Friston, 2005; Rao & Ballard, 1999; Wacongne et al., 2011). Given these varying theoretical perspectives, there are different views of what “prediction” is. For example, if prediction is based on pre-activation, anticipatory effects (e.g. evidence that the listener is actively considering a word as a perceptual hypothesis before any acoustic evidence for that word has been heard) can be taken as the litmus test of predictive processing. From a Bayesian perspective, however, processing that is not strictly anticipatory (i.e., where prior knowledge modulates processing of a word as it is being heard) is still predictive (Norris et al., 2016). There is also debate about whether prediction is the sole process underlying language comprehension, or whether it is only one of multiple processes (Huettig & Mani, 2016; Ito, Martin, & Nieuwland, 2017). Relatedly, there is current discussion about whether a key demonstration of prediction replicates (DeLong et al., 2005; Ito et al., 2017; Nieuwland et al., 2018). In spite of these ongoing debates, there is consensus that, at least under some circumstances (e.g. when the previous context is highly constraining), comprehenders can use contextual information to anticipate upcoming words and their associated features (Altmann & Kamide, 1999; Rommers, Meyer, Praamstra, & Huettig, 2013). This anticipatory ability is what we mean here by “prediction”.

Are visual and sentential cues therefore used jointly to predict acoustic-phonetic aspects of spoken words? At first glance, the electrophysiological data might suggest that prediction based on visual information is distinct from that based on sentential information: the N1 is several hundred milliseconds earlier than the N400. The timing of the N400 is such that N400 effects based on sentence context may not even reflect prediction: They are late enough to reflect instead effects of integration (where the current word is being integrated into the ongoing interpretation of the sentence) (Hagoort, 2008). If, however, visual and sentential constraints influence the same anticipatory process, we should expect to see effects of sentential context in the same time window as the N1. This hypothesis is supported by other evidence indicating anticipatory effects of sentential constraints on speech comprehension. In visual-world eye-tracking studies, for example, verb-based constraints about the type of noun that is likely to be the grammatical object of the verb can influence processing before the acoustic onset of the noun (Altmann & Kamide, 1999). Since sentential constraints can thus be used anticipatorily, and predictive processes based on visual information can be detected as changes to the N1, it ought to be possible to observe a modulation of the N1 visual effect by semantic constraints.

To test this potential early modulation, we used magnetoencephalography (MEG) to look at auditory N1 latency and amplitude in an audiovisual speech paradigm. The final words of spoken sentences could be contextually constrained or unconstrained and they began with salient or non-salient visemes. We predicted that the viseme effect (more N1 suppression to salient visemes) would depend on whether the sentence was constrained or not. Such a demonstration would suggest that sentential constraints are being used to predict (i.e., to anticipate) rather than to influence semantic integration. This is because the N1 is not considered to reveal semantic integration and because modulation of the N1 by viseme salience is a signature of form-based processing (i.e., it reflects predictions about the sentence-final word’s initial consonant, not its meaning). Such an interaction would thus suggest that effects of visual and sentential information in audiovisual speech comprehension are indeed two sides of the same coin, that is, that they both reflect the ability of the listener to predict upcoming words. We expected an interaction between viseme salience and sentential constraint to arise in the left hemisphere. We did not have a specific hypothesis about the shape of the interaction (whether a viseme effect would be prominent only in the absence of sentential constraints, or whether sentence and viseme would jointly suppress the neural response and shorten N1 latency). We also included the right hemisphere, where we did not expect to find an interaction, as a control.

## 2.1 Materials and Methods

### 2.1.1 Participants

Twenty-five Dutch native speakers participated in the experiment after signing an informed consent form in accordance with the Declaration of Helsinki. One was excluded due to excessive artifacts caused by makeup residue that had been magnetized during a magnetic resonance imaging (MRI) session earlier that day. None of the remaining 24 participants (6 male, aged 19-28, M = 22.25, SD = 2.17) reported neuropsychiatric disorders. All were right-handed, with normal hearing and normal or corrected-to-normal vision. The study was approved by the local ethics committee (CMO region Arnhem/Nijmegen), and participants were paid for their participation.

### 2.1.2 Experimental materials, design and procedure

Participants watched videos of a speaking person who was recorded from the shoulders up (see Fig. 1A). The video contained spoken sentences. The last word (target word) of the sentence either started with a salient or a non-salient viseme. A salient viseme allowed the upcoming auditory content of the target word to be predictable (relative to a non-salient viseme). The content of the target words could also be predictable depending on the form and meaning of the preceding sentence. We therefore orthogonally manipulated the viseme salience of the initial sounds of sentence-final words and the sentential constraints on those words (see Table 1). This resulted in four experimental conditions: constraining sentential context with salient viseme, constraining sentential context with non-salient viseme, unconstraining sentential context with salient viseme, and unconstraining sentential context with non-salient viseme. Additionally, we had the speaker say the target words in isolation to test for the early N1 effect of viseme salience without a sentence context. Here, our manipulation resulted in two experimental conditions: salient viseme and non-salient viseme.

**Figure 1.**
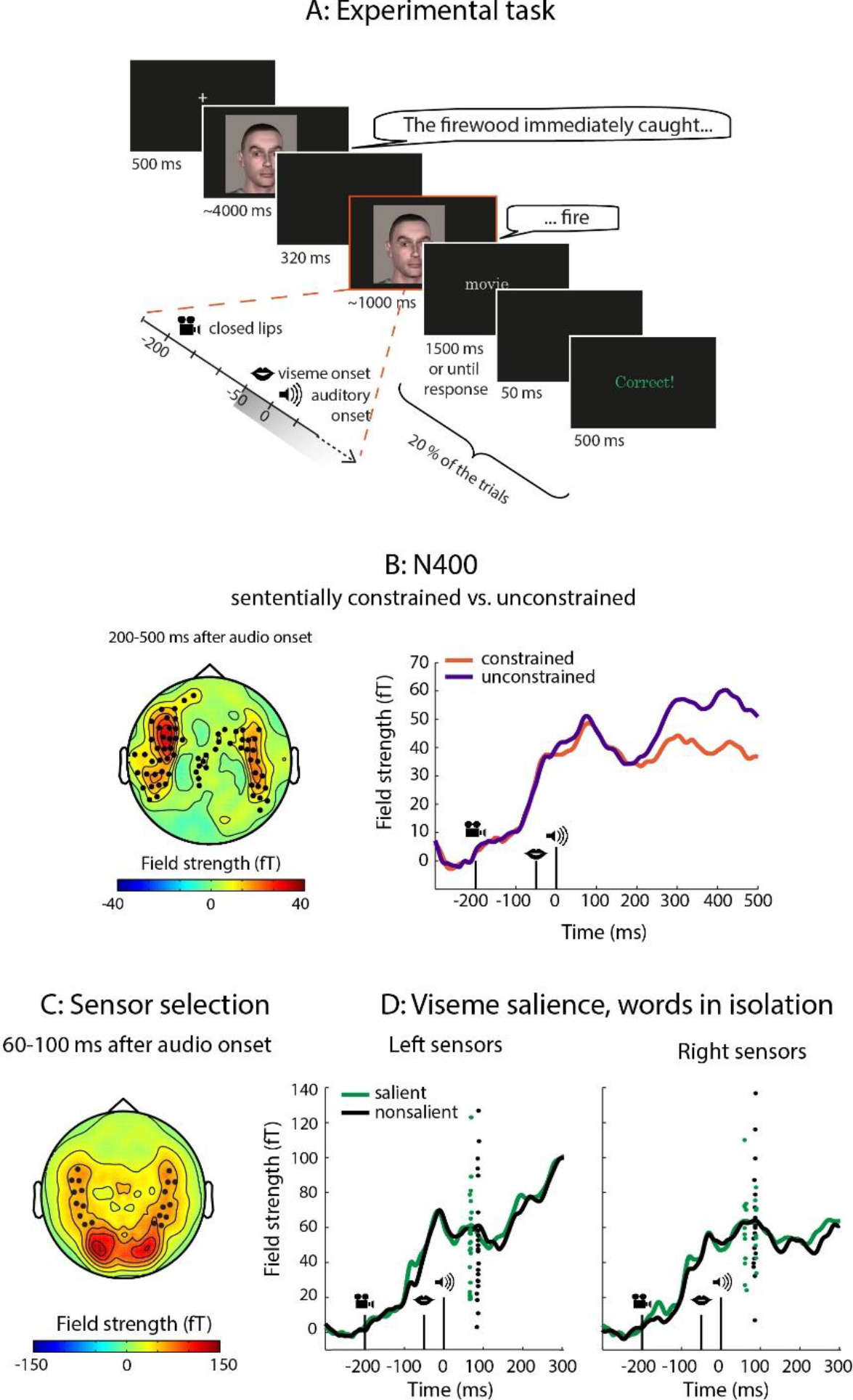
A: Trial sequence. A video displays a speaker uttering a Dutch sentence for about four seconds. The final word of the sentence appears after a short blank screen (320ms). In this example, the auditory contents of the sentence-final target word are made more predictable due to the salient viseme (/f/), and the constraining sentential context. The target word here (‘fire’) is followed by a written word, presented on 20% of the trials. Participants pressed a button to indicate whether it was the same as the previous word. B: N400-effect of sentence context. Left: topography of the difference in activity to words preceded by a sententially unconstrained vs. constrained context, with significant sensors highlighted. Sententially unconstrained words led to stronger neural activity over temporal, parietal and frontal sensors. Right: event-related fields to sententially unconstrained (purple) and constrained sentences (orange), averaged over sensors highlighted on the left. C: Sensors of interest with average N1 topography to videos of single words. The most active left and right temporal sensors are highlighted. D: Event-related fields to words beginning with salient (green line) and non-salient visemes (black line), plotted separately for sensors in the left and right hemisphere. Jackknifed auditory N1 peak latencies for each subject are represented by dots. In the left sensors, the auditory N1 peaked earlier if the viseme was salient than if it was not.

**Table 1.**
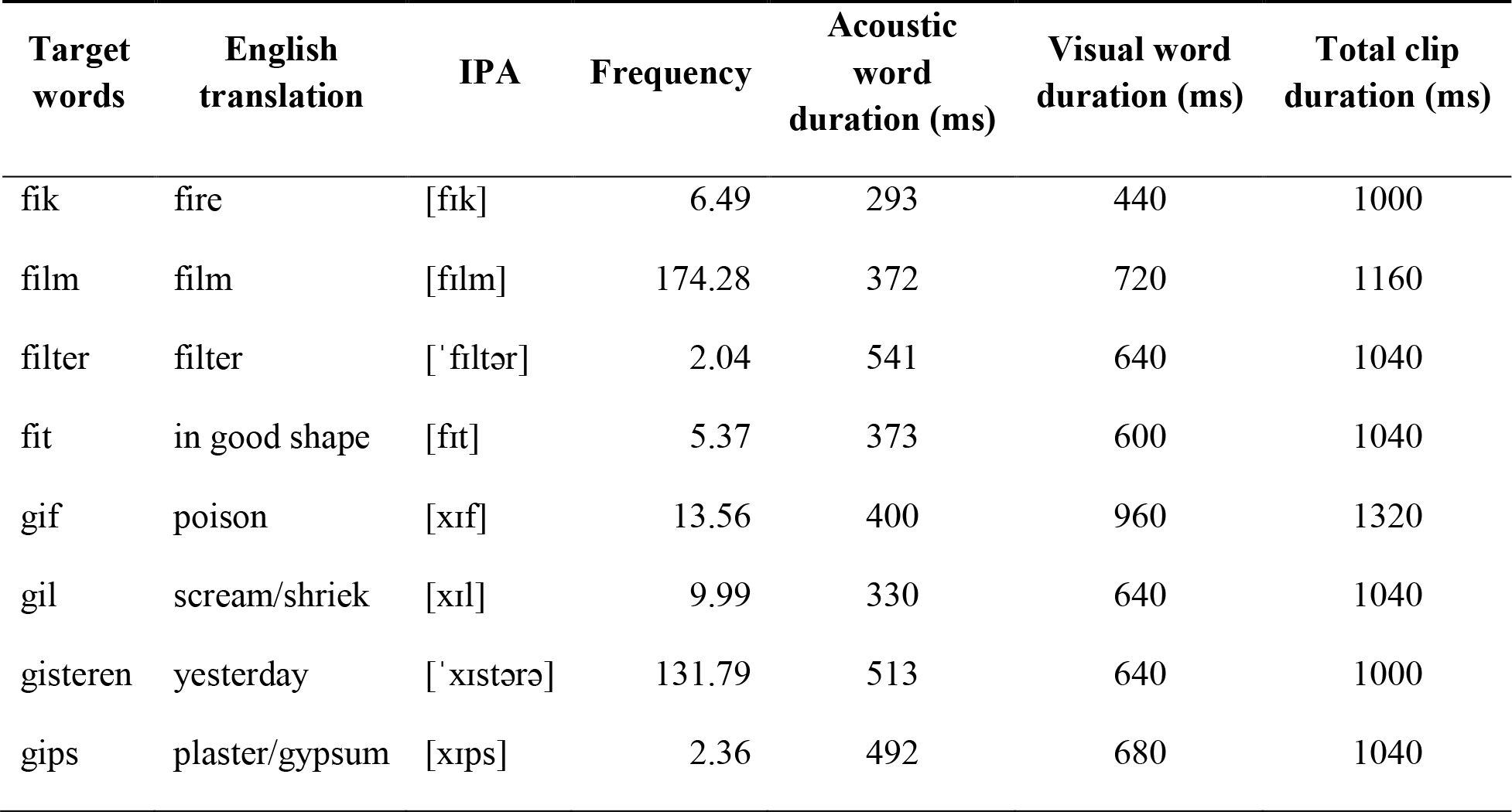
Target words. IPA = International Phonetic Alphabet. Frequency did not differ significantly between f- and g-words.

| Target words | English translation | IPA | Frequency | Acoustic word duration (ms) | Visual word duration (ms) | Total clip duration (ms) |
| --- | --- | --- | --- | --- | --- | --- |
| fik | fire | [fɪk] | 6.49 | 293 | 440 | 1000 |
| film | film | [fɪlm] | 174.28 | 372 | 720 | 1160 |
| filter | filter | [ˈfɪltər] | 2.04 | 541 | 640 | 1040 |
| fit | in good shape | [fɪt] | 5.37 | 373 | 600 | 1040 |
| gif | poison | [xɪf] | 13.56 | 400 | 960 | 1320 |
| gil | scream/shriek | [xɪl] | 9.99 | 330 | 640 | 1040 |
| gisteren | yesterday | [ˈxɪstərə] | 131.79 | 513 | 640 | 1000 |
| gips | plaster/gypsum | [xɪps] | 2.36 | 492 | 680 | 1040 |

#### Target words

We manipulated viseme salience while keeping acoustic features constant across and within conditions. In order to achieve this, we selected two Dutch speech sounds that are phonetically similar but that differ in visual salience: the unvoiced labiodental fricative /f/ (the same sound as an English f) and the unvoiced velar fricative /x/ (similar to the final sound in ‘*Bach*’). Fricatives are consonants we produce by forcing air through a narrow constriction. In the case of an /f/, this constriction is between the upper teeth and lower lip, making it visually salient. For /x/, the constriction is created by positioning the back of the tongue close to the soft palate, resulting in a less salient viseme. Based on auditory information alone, Dutch participants can correctly distinguish /f/ and /x/ after having heard approximately equal portions of the sounds (Smits, Warner, McQueen, & Cutler, 2003). In addition, these two sounds are often confused with each other early in the auditory recognition process (Smits et al., 2003). They are also similar in duration. To confirm this, we measured the frication duration for 7 /f/ and 7 /x/ words, yielding means of 143 ms and 147 ms respectively. Since /f/ and /x/ are acoustically similar, additional visual information could make their discrimination easier.

We then chose four words starting with each of these two speech sounds followed by the vowel /ɪ/ to further ensure acoustic similarity. This resulted in eight target words (Table 1). The mean frequencies of the words in the two conditions were not significantly different (t(6) = 0.145, p = 0.89) based on the word frequencies in the SUBTLEX-NL database (Keuleers, Brysbaert, & New, 2010). We visually inspected the sound clips in Praat (Oostenveld, Fries, Maris, & Schoffelen, 2011; Boersma, 2001) to determine acoustic duration. We determined visual duration (onset to offset of lip-movements) by visual inspection of the video clips in Adobe Premiere Pro 6. To minimize the effect of word repetition and strategic processing related to the onset sounds, we additionally introduced trials ending with 16 filler words, none of which started with /f/ or /x/. The filler words were semantically related but acoustically distinct from the target words (e.g., “verband” (bandage) and “klei” (clay) for the target word “gips” (plaster)). We did not analyse neural activity related to these filler trials.

#### Sentences

We initially constructed 24 different sentences ending with each of the eight target words, which makes a total of 192 sentences. Half of the sentences strongly predicted a particular sentence-final target word, whereas the other half were not strongly constraining, and thus poor predictors of the target words. We pilot tested how predictive the sentences were (without the final words) on a group of 40 Dutch participants (native speakers with no dyslexia) with one of four versions of a pen-and-paper sentence completion test. Participants were asked to indicate which word best completed each sentence. Based on the results of this test, we then selected the 10 most and 10 least reliably predictive sentences for each target word, which gave us a final set of 160 target sentences (see Appendix for all sentences and Table 2 for examples). The predictive power of the sentences selected for the experiment was on average 80% predictability of the sentence-final word. We also constructed 10 sentences for each of the sentence-final filler words, half of which were predictive and half of which were not.

**Table 2.** The four experimental conditions with example sentences.

|  | Salient viseme | Non-salient viseme |
| --- | --- | --- |
| **Sententially constrained** | Het brandhout vloog meteen in de FIK<br>*The firewood immediately caught FIRE* | De tanden van een cobra bevatten dodelijk GIF<br>*The teeth of a cobra contain deadly POISON* |
| **Sententially unconstrained** | Toen Roel thuiskwam stonden zijn spullen in de FIK<br>*When Roel came home his stuff was on FIRE* | De biologiestudenten lezen een artikel over GIF<br>*The biology students read an article about POISON* |

#### Video recording and editing

A male native Dutch speaker with a neutral dialect was chosen as speaker. He was seated in a chair facing the camera. A soft box lighting device was behind the camera to ensure maximal visibility of his facial movements. The speaker was instructed to speak clearly and at a natural pace, but to include a pause before the sentence-final word. Later, a single target word video was chosen and presented at the end of all 20 sentences associated with that word. The recording was done in a soundproof room using a digital HD video camera (JVC HD GY-HM100E) at 1920×1080 resolution and 25 progressive frames per second. Sound was recorded with the camera microphone at a sampling rate of 48 kHz.

First, video clips of each sentence and each sentence-final word were selected and cut out from the raw recordings in Adobe Premiere Pro 6. Every sentence and word video started and ended with a neutral face and a closed mouth. The video clips of the lead-in sentences had six frames (240ms) of neutral face at the beginning and three frames (120ms) at the end, whereas the word video clips started with three (120ms) and ended with six frames (240ms) of neutral face. This enabled us to control the visual gap between lead-in sentence and target word. The sound was noise-reduced in Audacity and its amplitude was root mean square equalized in Praat. We put the edited sound clips back into the video clips in Adobe Premiere Pro 6 without any audiovisual asynchrony or realignment, after which we exported the edited video clips in uncompressed AVI format with a 720×480 frame size. Finally, we compressed and exported the video clips as AVI files in the Indeo 5.10 codec using VirtualDub.

#### Stimulus presentation

Each trial started with a fixation cross placed approximately between the eyes of the speaker in the upcoming video. After 1500 ms, the fixation cross was replaced by the lead-in sentence video (in sentence blocks) or target-word video (in word blocks). In the sentence blocks there was a gap during which a uniformly black screen was presented for 320 ms between the lead-in sentence and the sentence-final target word (Fig. 1A). This ensured a constant interval (560 ms) between visual lead-in sentence offset and target word onset. Additionally, it allowed us to use identical target word videos across trials. During sentence blocks, a target word was never repeated on two consecutive trials, and the proportion of fillers and targets was balanced across blocks. In word-only trials the target and filler words were presented in a pseudorandom order. Importantly, the onset of the auditory content in the target word videos naturally lagged the viseme onset by 50 ms, and was constant within and across conditions.

### 2.1.3 Task

While participants performed the task, we recorded their ongoing neural activity with MEG. The video was projected on a screen at a distance of 70 cm from the participant, and the sound delivered binaurally through MEG-compatible air tubes at a comfortable sound pressure level. Stimulus presentation was controlled by a PC running Presentation software (Neurobehavioral Systems). There were 8 sentence blocks of 40 trials, leading to a total of 160 filler trials and 160 target trials. After the sentence blocks, there was one word-only block with 20 trials per target word, again leading to a total of 160 trials. The inter-trial interval varied randomly between one and two seconds throughout the experiment. All participants saw the same set of stimuli, but the order of presentation differed per participant.

To ensure that participants would pay attention to the target words, we added a word discrimination task on 20% of the sentence trials and 35% of the word trials. In these trials, a written word appeared on the screen immediately after the word video, and the participants had to indicate with a button press whether this word was the same or different from the word they had just heard.

Before the experiment began, participants tried two practice trials. During the experiment, there was a break of at least 30 seconds after every block of 40 sentence trials, and a break of at least 60 seconds before the word-only block. After these obligatory breaks were over, participants could press a button to proceed with the experiment whenever they wanted.

#### MEG acquisition

We recorded brain activity using a whole-head MEG (275 axial gradiometers, VSM/CTF Systems) at a sampling rate of 1200 Hz in a magnetically shielded room. Participants’ head position was monitored during the experiment using coils placed at the nasion and in both ear canals, and was corrected during breaks if needed. Horizontal and vertical electro-oculogram (EOG) was recorded using 10-mm-diameter Ag–AgCl surface electrodes. We later used the vertical EOG to aid offline rejection of blink artifacts.

### 2.1.4 MEG data analysis

The MEG data were preprocessed and analysed in Matlab (MathWorks, Natick) using the FieldTrip toolbox (Oostenveld, Fries, Maris, & Schoffelen, 2011). We calculated event-related fields (ERFs) time-locked to the onset of the target word videos, and used these to analyse auditory N1 latencies as well as early and late amplitude effects as a function of our experimental manipulations.

#### Calculation of event-related fields

We extracted trials of 1300 ms from the data starting 300 ms before the onset of the target word videos. Trials containing jumps in the MEG signal caused by the SQUID electronics were rejected based on visual inspection. Trials containing excessive muscle artifacts were then rejected after visual inspection of the amount of variance in an epoch with the signal bandpassed at 110-140 Hz. Next, we used independent component analysis (Bell & Sejnowski, 1995) to remove variance in the signal pertaining to eye blinks (Jung et al., 2000). Finally, we discarded any remaining trials where the ERF amplitude was more than 4 standard deviations above or below the mean. In total, we rejected an average of 12.75 (SD = 4.80) trials per participant.

Before calculating ERFs for each condition of interest, we low-pass filtered the data at 40 Hz and baseline corrected relative to a 100 ms time window before the onset of the word videos. Finally, we calculated planar gradient transformed ERFs (Bastiaansen & Knösche, 2000), a procedure which simplifies the interpretation of the sensor-level data because it places the maximal signal above its source (Hämäläinen, Hari, Ilmoniemi, Knuutila, & Lounasmaa, 1993). Importantly, this operation removes the polarity of ERF components, making the strength of their deflections from zero across conditions the main information of interest.

#### Sensors of interest

We constrained our analyses to temporal sensors, with the exclusion of sensors bordering occipital areas (in order to avoid excessive contamination by visual activity). To further select the sensors of interest, we used the grand averaged ERF data from words presented in isolation. We selected the 10 most active temporal sensors in each hemisphere in a time window corresponding to the auditory N1 component, 60-100 ms after auditory onset (Fig. 1C). We have used the same sensor selection procedure for all previous auditory studies in our lab.

#### Analysis of auditory N1 peak latency

We calculated auditory N1 peak latencies for the target words, separately for each hemisphere. We used a jackknife approach, which allows for robust estimation of latency differences, to estimate the N1 peak latencies (Kiesel, Miller, Jolicœur, & Brisson, 2008; J. Miller, Patterson, & Ulrich, 1998). Instead of computing one average N1 latency value per participant, we computed as many averages as there are participants while leaving one participant out each time. If the latency is consistent over participants, then the average value will not change substantially depending on which participant is left out. To test whether the conditions of interest exhibited latency differences, we compared the estimated peak latencies for salient vs. non-salient visemes in the time window between 60 and 100 ms after auditory onset, with the t-values corrected to t_corr_ected = t / (n-1) in order to reduce the false positive error rate (Ulrich & Miller, 2001). We first compared latencies in the word-only condition, then for words in sentence contexts. To assess whether viseme salience modulations of N1 latency depend on sentential predictability, we also tested the viseme/context interaction by comparing the difference of the viseme effects in the two sentential conditions against each other.

#### Analysis of auditory N1 amplitude

We tested for auditory N1 amplitude differences as a function of the preceding viseme and sentential information. We used nonparametric cluster-based permutation t-tests for paired samples (Maris & Oostenveld, 2007). Cluster-based permutation tests are well suited to test for differences between conditions in time-series data. The test controls for the false alarm rate by taking advantage of the fact that effects are typically clustered in time. We tested for clusters of amplitude differences across samples of a 300 ms long time-window starting from 100 ms before auditory onset (i.e., 50 ms before viseme onset, 100-400 ms after target word video onset,), using 5000 permutations for the generation of the null distribution. We first compared salient to non-salient visemes in the word-only condition, followed by the same comparison for words in a sentence context. We also compared the effect of constraining vs. unconstraining sentential contexts, and, finally, we tested the viseme/context interaction. All reported amplitude p-values are cluster p-values.

#### N400 analysis

To verify that our sentence context manipulation resulted in a classical N400 effect (Kutas & Federmeier, 2011), we used a cluster test to compare ERFs of the sententially constrained and unconstrained sentence-final words. This test searches for groups of sensors that show a significant difference between the two conditions. We collapsed over all time points from 200 ms to 500 ms after auditory onset for this analysis.

## 3.1 Results

### 3.1.1 Task accuracy

On 20% of the trials, participants performed the word discrimination task. They did so with high accuracy (95% + 5.78%, mean + SD) suggesting that they paid attention to the target words.

### 3.1.2 N400 amplitude decreases in constrained sentence contexts

We first examined whether our manipulation of sentence context resulted in an N400 effect. The N400 component to a sentence-final word is known to decrease in amplitude in the presence of a constraining context, that is, when the sentence context renders it predictable (Kutas & Federmeier, 2011). This is what we found as well: a large number of sensors showed an attenuated response in the constraining sentential context compared to the unconstraining sentential context (p = 0.015, pre-defined time window of 200-500 ms after auditory onset). The difference topography in this time window (Figure 1B) suggests that the effect of sentential constraints was present in both hemispheres, but was more pronounced on the left side. It originated from a difference in activity in temporal sensors, as well as a smaller number of parietal and frontal sensors.

### 3.1.3 Salient visemes shorten auditory N1 latency in the left hemisphere

We assessed whether viseme salience influenced the peak latency of the auditory N1 to words presented in isolation, outside of a sentence context (Fig 1D). We were able to extract a reliable N1 peak latency for both viseme conditions (salient and non-salient) in the left hemisphere. In contrast, in the right hemisphere, the jackknife-estimated peak latencies following salient visemes formed a bimodal distribution, indicating two separate peaks of similar amplitude (i.e. an absence of a reliable N1 peak). We therefore compared the peak latencies only in the left hemisphere. There we found that words beginning with salient visemes were associated with an earlier auditory N1 peak than words beginning with non-salient visemes (t (23) = 2.25, p = 0.034).

We next tested whether the viseme effect - earlier N1 peaks to words containing salient compared to non-salient visemes - would continue to be present when the words were embedded in a (constraining or unconstraining) sentence context (Fig 2C). We performed this comparison in the left hemisphere only. We found that salient visemes shortened the N1 latency in an unconstrained sentential context, in other words when the upcoming word could not be predicted (mean difference = 21 ms, t (23) = 2.15, p = 0.02). We did not find an effect of viseme salience in the constrained sentential context (mean difference = 6 ms, t (23) = 0.98, p = 0.41). However, we did not find evidence for an interaction effect between viseme salience and sentential context (no difference in the viseme effect in constraining compared to unconstraining sentences, mean difference of differences = 15 ms, t (23) = 1.49, p = 0.147). In sum, although the viseme effect persisted when words were embedded in unconstraining sentences, the lack of an interaction effect makes it difficult to estimate the extent to which sentential constraints do or do not influence how much viseme salience influences auditory N1 peak latency.

**Figure 2.**
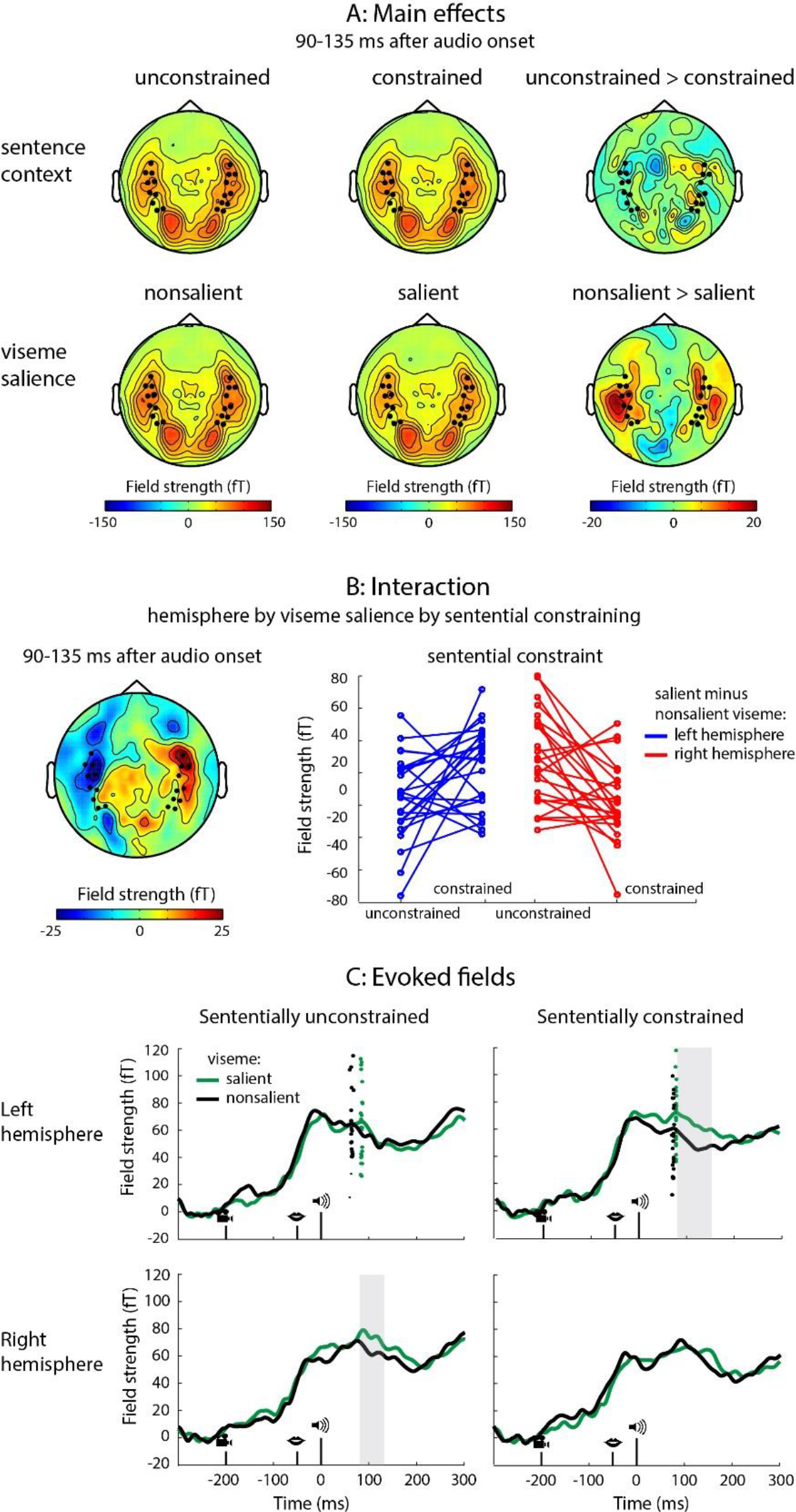
A: Topography of the main effects of sentential constraints and viseme salience in the time window of the significant three-way interaction between hemisphere, viseme salience and sentence context. Sensors of interest are highlighted. B: Left - Topography of the interaction. Right - Individual subject representation of the interaction, in two most prominent sensors on each side. Each red and blue dot represents the difference in the signal between non-salient and salient visemes, under different conditions of sentential constraint. C: ERFs for the salient and non-salient visemes in the two hemispheres under different conditions of sentential constraint. Clusters of significant differences are highlighted. Dots in the upper plots (left hemisphere) represent individual jackknife-estimated N1 latencies.

### 3.1.4 The early joint influence of viseme salience and sentential constraints is hemisphere-dependent

We looked at ERFs (up to 300 ms after word onset) to assess whether viseme salience, which exerted a consistent influence on N1 latency in the left (but not right) hemisphere, would also exert an effect on signal amplitude. When words were presented in isolation neither the left (p=0.137) nor the right (p=0.131) hemisphere showed evidence of an amplitude difference in early auditory processing following visemes of different salience (Fig 1C).

We then asked whether viseme salience affects signal amplitude to words presented in the context of a sentence. We found that viseme salience and sentential constraints came together to exert a joint influence on auditory processing, but that the pattern of their interaction depended on the hemisphere (Fig 2 B, C). In both hemispheres, non-salient visemes led to larger auditory responses in the N1 time window. However, this viseme effect was modulated by sentential constraints in opposite ways in the two hemispheres. A viseme effect was evident only in the *presence* of sentential constraints in the left hemisphere, and only in the *absence* of sentential constraints in the right hemisphere (three-way interaction between hemisphere, sentential constraints and viseme salience: p=0.012, 90-135 ms). In other words, the left auditory cortices responded less to salient visemes when the upcoming word could be predicted based on sentential constraining while the right auditory cortices responded less to salient visemes when the upcoming word could *not* be predicted using sentential constraints (Fig 2B, left).

Next, we looked at the effect of viseme salience and sentence context for the two hemispheres separately. In the left hemisphere, we found no evidence for an effect of sentential constraints on early auditory responses (main effect of sentential constraints: no clusters found). Viseme salience, in contrast, did exert an effect on the early neural response (main effect of viseme salience: p=0.012, 80-126 ms after sound onset). The stronger amplitude to non-salient than salient visemes depended on whether a sentence was sententially constraining or not (marginal interaction between viseme salience and sentence context: p=0.061, 131-159 ms). Namely, a viseme effect was present if the sentences were sententially constrained (p=0.04, 71-152 ms), but we found no evidence of it if the sentences were sententially unconstrained (p=0.265).

In the right hemisphere, we observed a reverse pattern of results. We found no evidence of an effect of viseme salience on neural activity (no main effect of viseme: p=0.196). Conversely, we observed stronger activity to sentence-final words preceded by an unconstraining sentential context relative to a constraining one (main effect of sentential constraint: p=0.020, 12-51 ms). As in the left hemisphere, in the right hemisphere we also found a combined effect of whether a viseme was salient and whether the sentence was sententially constrained (interaction between viseme salience and sentential constraint: p=0.044, 87-117 ms). Here, words starting with non-salient visemes led to more neural activity than words starting with salient visemes in sententially unconstraining sentences (p=0.040, 90-121 ms), but we found no evidence of a viseme effect in sententially constraining ones (p=0.278).

## 4. Discussion

In this study, we examined the independent and joint effects of viseme salience and sentential constraints on early auditory processing in sentence-final words. In electrophysiological studies, viseme salience typically affects N1 latency and/or amplitude (Baart, 2016), while sentential constraints typically affect the N400 (Kutas & Federmeier, 2011; Kutas & Hillyard, 1980; Lau et al., 2008; Maess, Herrmann, Hahne, Nakamura, & Friederici, 2006). If, however, both effects reflect an increase in the predictability of the upcoming auditory input, and since studies using other paradigms (e.g. eye tracking) (Altmann & Kamide, 1999), have demonstrated effects of sentential context even earlier than the N1, we predicted that sentential constraints could modulate the early viseme effect by making the viseme more or less predictable. We found that sentential context can indeed have an effect on early language processing (~90 ms after word onset) through modulating reliance on early visual cues, but that the pattern of this effect depended on the brain hemisphere: the left hemisphere integrated probabilistic cues from visual and sentential information, while the right hemisphere gave visual cues priority only when no strong sentential constraints were present.

Before we go on to discuss these results in greater detail, it is important to note where our study stands in terms of ecological validity. The majority of studies on viseme salience investigate the effect in isolated phonemes or syllables (Arnal et al., 2009, 2011; Baart, 2016; Wassenhove et al., 2005). In contrast, we employed a paradigm where full sentences were spoken. However, we introduced a break between the penultimate word and the target word. This manipulation allowed for better experimental control and a higher signal to noise ratio, but it comes at the cost of reducing predictability based on natural prosody and coarticulation. While we believe that our results allow for the conclusion that comprehenders *can* use both viseme salience and sentential constraints to predict upcoming words, they do not necessarily imply that people always do so in natural speech situations. In addition, we repeated the target words throughout the experiment. This might have increased anticipation of their identity in all experimental conditions, which could have led to an attenuated neural signal based on both higher predictability and repetition suppression, and therefore smaller observed differences between the conditions. In addition, as the target words were task-relevant, the increased predictability might have also increased attention to them. It remains an open question whether the pattern of observed results would hold with task-irrelevant words.

We first replicated a typical N400 effect (a decreased amplitude to words embedded in a constrained compared to an unconstrained sentence context, Fig 1B). This suggests that participants were anticipating upcoming word forms when the sentential context was constrained, although it might equally reflect ease of integration (Hagoort, 2008). We also replicated a viseme effect (shorter N1 latencies to words beginning with a salient viseme compared to a non-salient one) in left temporal sensors, when words were presented outside the context of a sentence, however we did not find an effect on the N1 amplitude. The latency shortening implies that participants used the visual information from the lip movements to predict upcoming auditory input. This type of latency shortening has been argued to reflect faster auditory processing (Wassenhove et al., 2005), and to be insensitive to whether or not the speech sound matches the viseme (Arnal et al., 2009). One suggested mechanism underlying visual facilitation of auditory speech processing is phase-resetting of activity in auditory cortex due to input from visual motion areas (Arnal et al., 2009; Schroeder, Lakatos, Kajikawa, Partan, & Puce, 2008).

Crucially, when we investigated auditory activity to words embedded in the context of a sentence, we found that the level of sentential constraints modulated both the N1 latency and its amplitude, through changes in the effect of viseme salience. Both hemispheres exhibited joint sensitivity to viseme salience and sentential constraints, but, interestingly, the effect of these two factors on early auditory word processing were differently expressed. In the left hemisphere, we observed N1 peak latency shortening to salient visemes in the absence of sentential constraints, as well as an amplitude reduction in the presence of sentential constraints. In the right hemisphere, we did not detect reliable N1 peaks (and so did not estimate latencies or a possible hemispheric interaction), but a lower amplitude to salient visemes was evident in the absence of sentential constraints.

A number of studies have demonstrated a suppressed and earlier N1 for words beginning with more salient visemes (Baart, 2016; Klucharev et al., 2003; Wassenhove et al., 2005), and one study suggested that the degree of salience could be predictive of the degree of suppression of the BOLD response (Arnal et al., 2011). We replicated this stronger suppression to words beginning with more salient visemes in the left hemisphere when prior sentence context was constrained, but not when it was unconstrained. In contrast, the N1 *latency* to salient visemes in the left hemisphere was shorter only in the absence of sentential constraints. It is important to note that, even though we found an N1 peak latency shift to salient visemes when the target words were not sententially constrained, and no viseme effect when they were, we did not find evidence of a viseme-context interaction in this analysis. This is partially in line with a recent EEG study that combined sentential constraints with viseme salience, where there was an N1 latency shift only for salient (vs. non-salient) visemes, but no early interaction with sentential constraints (Brunellière et al., 2013).

We also looked at the effect of viseme salience and sentence context on the ERF amplitudes. Both hemispheres were sensitive to a combination of sentential constraints and viseme salience in the N1 time window, but surprisingly, the reliance on viseme salience as a function of sentential constraints differed per hemisphere, with the left hemisphere integrating both effects, but the right hemisphere giving priority to visual cues only in the absence of sentential constraints. In the right hemisphere, we found a higher amplitude for non-salient visemes in the absence of sentential constraints, indicating a reliance on visual cues only when there was no helpful sentential context. In the left hemisphere, conversely, the combined effect of sentential and visual cues suppressed neural activity jointly. Here it appears that the presence of sentential context rendered the word form, and hence the viseme itself, predictable, thus facilitating subsequent auditory processing. The resulting topography of this early hemisphere by context by salience interaction is bilateral and symmetric (Fig. 2B, left).

Why would the joint effect of viseme salience and sentence context (both of which make an upcoming word more predictable) lead to a reduction in the early auditory neural response to words in the left hemisphere? Recent research proposes a major role for stimulus likelihood. Namely, expecting an upcoming auditory stimulus attenuates the auditory N1 both for pure tones (Hughes, Desantis, & Waszak, 2013; Lange, 2009; Todorovic et al., 2011; Todorovic & Lange, 2012) and for spoken words (Houde, Nagarajan, Sekihara, & Merzenich, 2002). Auditory predictions related specifically to viseme salience also cause this early attenuation (Arnal et al., 2009, 2011).

Where in the brain do these predictabilities exert their influence? The topographies of our early effects suggest a broad bilateral effect in the temporal lobes for viseme salience (Fig. 2A), and a slightly more anterior, more constrained effect for the interaction between viseme salience and sentential context (Fig. 2B). Broad activity along the superior temporal sulcus has previously been found both when people observed lip movements without sound and when they heard speech (Skipper, van Wassenhove, Nusbaum, & Small, 2007). The sensitivity to mismatch between visual and auditory speech has also been suggested to rely on a feedback signal from the superior temporal sulcus (Arnal et al., 2009). In addition, the anterior temporal cortex has been implicated in several studies of sentential processing (Humphries, Love, Swinney, & Hickok, 2005; Vandenberghe, Nobre, & Price, 2002), and we find it a likely candidate for the source of our interaction effect. In fact, in one study activity related to the predictability of semantic priming was localized in the left anterior superior temporal gyrus (Lau, Weber, Gramfort, Hämäläinen, & Kuperberg, 2016), indicating a sensitivity to probabilistic semantic processing. In other words, our interaction of probabilistic processing appears more closely associated with areas corresponding to semantic processing than to those related to viseme processing alone.

A striking finding in this study is that both cortical hemispheres displayed an early sensitivity to viseme salience and sentential context, but that the pattern of this sensitivity differed. We did not expect to find a hemispheric interaction; however, the topography of the interaction appears convincingly symmetric, suggesting that it is constrained to the same brain area. This adds to a growing body of research that demonstrates an organization of language processing within the ventral stream which is bilateral, but with a hemispheric asymmetry in activation (Hickok & Poeppel, 2007; Stefanatos, Gershkoff, & Madigan, 2005). Behavioural research also supports the idea that the left and right hemisphere both process visual information, but in slightly different manners. For example, it has been claimed that the right hemisphere relies on surface-type visual information longer, whereas the left hemisphere has quicker access to deeper levels of lexical representation: when participants are asked to recognize a letter in a visual stimulus, the right hemisphere, as opposed to the left hemisphere, displays no word superiority effect (Krueger, 1975). When people have to complete words based on the first few letters, the right hemisphere displays a stronger effect of priming by previously seen words if the case in which the prime was presented matches the case in which the beginning of the target word is presented (Marsolek, Kosslyn, & Squire, 1992). It also appears that the right hemisphere accesses semantic information later, and keeps semantic information active longer, with semantic priming showing effects with larger a prime-to-target stimulus onset asynchrony than in the left hemisphere (Abernethy & Coney, 1996; Anaki, Faust, & Kravetz, 1998; Burgess & Simpson, 1988; Chiarello, Burgess, Richards, & Pollock, 1990; Chiarello & Richards, 1992; Koivisto, 1997). Here we show that, while *probabilistic* language processing is also bilateral, the pattern of neural responses conforms to the functional specificity of the two hemispheres. In the right hemisphere, we found a viseme effect only in the absence of sentential constraints, indicating a stronger reliance on visual information. The left hemisphere, in contrast, appears able to combine the probabilistic information involved in sentential constraints and viseme salience.

## 5. Conclusion

Spoken language processing requires integration of information across time, and one of the means that comprehenders have at their disposal to achieve this is that they can make predictions about upcoming content based on preceding content. These predictions vary in strength, arise from different cues, and are made at different levels of language processing. It remains to be determined what the functional mechanism is (e.g., pre-activation, Bayesian inference, generative modelling or predictive coding), but at the neural level it appears that there is suppression of neural activity to predictable sounds and words. Our study sheds light on the joint effects of viseme salience and sentential constraints. We found that both of these factors have an effect on early auditory processing (in the N1 range). The two hemispheres however handled this combined information differently, with the right hemisphere giving priority to visual information in the absence of strong sentential constraints, and the left hemisphere combining visual and sentential information. This speaks to a complex hierarchy of predictions in language processing, one that is reliant on general probabilistic processing mechanisms but is simultaneously highly dependent on the functional specificity (and lateralization) of the associated cortical areas.

## Acknowledgments

We would like to thank Vedran Dronjić for his input on hemispheric differences in processing semantic information.

## 7. Appendix

### Stimulus materials – target words

highconstr = high constraining sentence context

lowconstr = low constraining sentence context

#### gif_lowconstr

De oude vrouw dronk per ongeluk een beker met gif

Het water was gemengd met een eetlepel gif

In de avond zag hij een oude documentaire over het gebruik van gif

In een scheikundeles sprak de docent over het gebruik van gif

De grote fabriek produceert te veel gif

De man was geïnteresseerd in een nieuw boek over gif

Het speelgoed dat de moeder voor haar kind had gekocht bevatte gif

De biologiestudenten lezen een artikel over gif

In het experiment gebruikten de onderzoekers grote hoeveelheden gif

Sommige bacteriën produceren gif

Hij voelde zich slecht na het innemen van het gif

Ze herinnerde zich de naam van het gif

#### gif_highconstr

Ze hebben de ratten gedood met gif

Socrates stierf aan een beker met gif

Sommige slangen wurgen hun prooi, andere doden hun prooi met gif

Vogelspinnen verlammen hun prooi door haar te injecteren met een dodelijk gif

Kikkers met felle kleuren zijn gevaarlijk om te eten, want hun huid bevat gif

Aboriginals dopen hun pijlen in verlammend gif

Een executie in de VS wordt meestal uitgevoerd door een injectie met gif

De tanden van een cobra bevatten dodelijk gif

De moordenaar besprenkelde het eten met een dodelijk gif

Sneeuwwitje stierf aan een appel die gedrenkt was in gif

Je wordt ziek van het eten van een vliegenzwam, want die bevat een soort gif

Sommige paddenstoelen kun je niet eten, want die bevatten gif

#### gisteren_lowconstr

De man zag de naam van zijn zusje in de deelnemerslijst van gisteren

De familie keek naar de leuke foto’s van gisteren

In de lunchpauze wilde ze dezelfde salade als gisteren

Hij denkt graag aan de gebeurtenis van gisteren

Een eerste poging om brood te bakken deed Anne gisteren

Toen hij erover nadacht was hij erg blij met de reactie van gisteren

Ze waren duidelijk beschaamd over het gesprek van gisteren

De journalist was bezig met het conceptartikel van gisteren

De vrouw kon zich niets herinneren van gisteren

De studenten werkten veel harder dan gisteren

Het biertje in de glas smaakt anders dan dat van gisteren

Ze denkt niet dat Peter ziek is want ze zag hem gisteren

#### gisteren_highconstr

Het meisje werd vandaag geboren, maar de bevalling begon gisteren

Hij was een dag te laat gekomen; de conferentie begon gisteren

De oude vrouw herinnerde zich haar bruiloft als de dag van gisteren

Hij herinnerde zich zijn terugkomst in Nederland als de dag van gisteren

Ruim veertig jaar later herinnert zij zich het incident nog als de dag van gisteren

Ze herinnert zich haar eerste schooldag als de dag van gisteren

Janne herinnert zich de geboorte van haar eerste kind als de dag van gisteren

De vakantiegangers hopen dat het vandaag mooier weer wordt dan gisteren

Hij was te laat, haar verjaardag was gisteren

De jongens praten in de pauze enthousiast over Martijns feestje van gisteren

Elise bracht restjes voor de lunch van het diner van gisteren

De student had in de gaten dat de professor loog; ze was niet van gisteren

#### gips_lowconstr

Op de tafel stond er een enorm figuur van gips

Het ontwerp was niet erg stabiel, want het was gemaakt van gips

In het pakhuis stond een grote stapel blokken van gips

Na een week als stagiair, leerde hij de samenstelling van gips

Het model van het huis was gemaakt van gips

De bemanning die bij het transport zou helpen vergat het standbeeld van gips

Het bedrijf waarbij ze hun voorraden bestelden had geen gips

Zijn functie was toezicht houden op de productie van gips

De vrachtwagen was geladen met een aantal ton gips

In de cursus leerden ze over het maken van gips

Ze had geen schoenen aan vanwege het gips

Veel van de fabrieken in het land produceerden ook gips

#### gips_highconstr

Het meisje viel van de trap en toen moest haar arm in het gips

Als je iets gebroken hebt, moet het in het gips

Gistermiddag brak Peter zijn been op de halfpipe en toen moest zijn been in het gips

Een dokter zet gebroken lichaamsdelen in het gips

De dierenarts zette de gebroken poot van de hond in het gips

De arm van de pechvogel zat in het gips

Voor een beugel maken tandartsen een mal van je gebit en daarin gieten ze gips

Na het ongeval met de fiets zat zijn arm in het gips

Tijdens het verbouwen maken we de muren glad met witte platen van gips

Bij het isoleren van een muur komt over de glaswol een witte plaat van gips

In Nederland bekleedt men soms de muren met witte platen van gips

Veel ouders maken afdrukken van de handjes en voetjes van hun kind in gips

#### gil_lowconstr

Terwijl ze aan het fietsen waren, hoorden ze een gil

De politieagent die bij de gebeurtenis aankwam hoorde een verdachte gil

De kinderen waren aan het spelen en Lisa gaf een gil

Terwijl ze thee zaten te drinken hoorde iemand een gil

Vanwege de harde muziek hoorde geen van de feestgangers de gil

Iedereen die in de tuin zat was er zeker van, het was een gil

Het laatste dat hij verwachtte te horen, was een gil

Lisa bleef werken en leek niet beïnvloed te worden door de gil

De treinreizigers hoorden opeens een gil

De deur was gesloten en daardoor hoorde niemand zijn gil

Elke dag op weg naar het werk hoort hij een gil

Toen ze in de rij stonden om te betalen, hoorden ze een gil

#### gil_highconstr

Toen ze schrok, gaf ze een harde gil

Bij het zien van de muis gaf ze een harde gil

Indien je vragen hebt, geef je maar een gil

Uit het donkere bos klonk ineens een hoge, harde gil

Zodra je klaar bent, geef je maar een gil

De vrouw in de horrorfilm was erg bang toen ze het lijk zag en gaf een gil

Ze sprong in het ijskoude water en gaf een harde gil

Als je iets wilt hebben, geef je maar een gil

De meisjes die in het spookhuis zaten schrokken en gaven een luide gil

Toen hij het spook zag gaf de man een harde gil

De twee vriendinnen die elkaar weer ontmoetten gaven allebei een hoge gil

Toen ze de spin onder de kast vandaan zag komen gaf ze een luide gil

#### fit_lowconstr

Na een avondje uit voelt hij zich zelden fit

De oude vrouw gaat iedere dag naar de winkel en is heel fit

Als je deze pil iedere dag neemt word je erg fit

Haar vriendin kijkt graag televisie, maar ze is erg fit

Vorig jaar ging ze vaak op stap en was niet heel fit

De man die vanuit de spiegel naar hem keek was erg fit

De vader en moeder van Dennis zijn allebei redelijk fit

Van werken op kantoor word je niet erg fit

Het jonge meisje met de bril bleek heel fit

De vrouw in de krant vandaag was ongewoon fit

De zes maanden zwangere vrouw voelde zich fit

De testresultaten waren onbetwistbaar, Simon was erg fit

#### fit_highconstr

Ondanks zijn gevorderde leeftijd was de man heel gezond en fit

Als je regelmatig sport, word je erg fit

Marije speelt elke dag squash en wordt daardoor erg fit

Een topatleet is meestal erg gezond en fit

Een week voor de marathon voelde ze zich nog niet helemaal fit

Ze was ontslagen uit het ziekenhuis, maar voelde zich nog niet helemaal gezond en fit

Van regelmatig baantjes trekken in het zwembad word je gezond en fit

Als je een goede conditie hebt, ben je erg fit

Haar broer is voetbaltrainer en erg fit

Als je elke dag een uurtje wandelt word je zeker fit

Iemand die snel buiten adem is tijdens het sporten, is niet erg fit

Tim volsnuit twee pakjes zakdoekjes per dag, en is niet erg fit

#### fik_lowconstr

De boeken van oma en opa stonden in de fik

Hun mooie tafel en stoelen stonden in de fik

De televisie van de jongen stond niet meer in de fik

Toen Roel thuiskwam stonden zijn spullen in de fik

Ze steken een aantal spullen uit het oude huis in de fik

Toen hij terugkwam van de winkel stond zijn auto in de fik

De gloednieuwe piano stond gelukkig niet in de fik

Het grootste deel van de meubels van het echtpaar stonden al in de fik

Het was moeilijk om overzicht te krijgen, want alles stond in de fik

Tot haar verbazing stonden haar schoenen ook in de fik

De antieke grammofoon uit de negentiende eeuw stond in de fik

Ze verzamelde al het papier en stak het in de fik

#### fik_highconstr

Met één lucifer stak hij de hele bovenverdieping in de fik

De pyromaan stak het huis in de fik

De tafeldecoratie stond te dicht bij het kaarsje en vloog plotseling in de fik

De brandweer kwam snel want het gebouw stond in de fik

Met Driekoningen staken we oude kerstbomen in de fik

Het brandhout vloog meteen in de fik

Hij had niet uitgekeken met zijn sigaret en zijn tent stond plotseling in de fik

Loek stak per ongeluk een stapel houten pallets in de fik

Een van de auto’s in het auto-ongeluk vloog ineens in de fik

De lekkende olie uit de frituurpan vloog ineens in de fik

Voor ze het wist, stond de hele stapel met aanmaakblokjes in de fik

Kijk uit met vuur, want voor je het weet staat alles in de fik

#### filter_lowconstr

Pim dacht dat er iets mis was met de filter

De onderzoekers goten de vloeistof door een filter

Hij kocht een nieuwe stofzuiger, maar er ontbrak een filter

Ze ging naar de winkel en kocht een nieuw filter

Als het systeem verstopt is, vervang je gewoon de filter

De installatie moest opnieuw gedaan worden vanwege een kapotte filter

Om de ruis te verwijderen gebruikten we een filter

De nieuwe machine zal niet werken zonder de filter

Ze moest kiezen tussen een rood en een groen filter

Veel apparaten die we thuis hebben bevatten een filter

Geen van de ingenieurs stellen vraagtekens bij de filter

Hij volgde de instructies op en goot de drank door de filter

#### filter_highconstr

Hij wilde jus d’orange zonder pulp maken en gebruikte daarvoor een pers met een filter

De oude man rookte nog altijd sigaretten zonder filter

Om schoon drinkwater te krijgen in landen waar het vervuild is, gebruikt men een filter

Het water in een aquarium wordt gezuiverd door een pomp met een filter

Om grondwater te veranderen in schoon drinkwater gebruik je een filter

In een zuiveringsinstallatie stroomt al het water door een pomp met een filter

Er loopt geen koffie meer door de trechter, vanwege de verstopte filter

We kunnen van zeewater drinkwater maken met behulp van een filter

Ze wilde het harde water zachter maken en gebruikte daarvoor een filter

Het laboratorium installeerde een luchtzuiveringssysteem met een filter

De zuiveringsinstallatie werkte niet vanwege een verstopte filter

Om koffiedik van de koffie te scheiden gebruik je een filter

#### film_lowconstr

Beide initiatiefnemers waren zeer tevreden met de film

Het enige dat haar vrolijk maakt wanneer ze verdrietig is, is een leuke film

Sommige van de kinderen kwamen helaas te laat voor de film

Het verjaardagscadeau van haar moeder was een film

Alex wachtte een heel jaar op de nieuwe film

Vorige week las ze een artikel over een oude film

Na het avondeten ging Frank door met lezen over de film

Ze maakt vaak haar laatste zakgeld op aan een film

Elke nacht droomt Miriam over dezelfde film

Het liedje herinnerde haar altijd aan een bepaalde film

Ze was erg overstuur na het zien van de film

Hij heeft een goed geheugen en herinnerde zich alle details van de film

#### film_highconstr

Op zaterdagavond kijkt ze vaak naar een leuke film

Will Smith is een van de acteurs in de nieuwe film

In de bioscoop is er vanavond een première, dus hij gaat vanavond naar de film

Het personeel van de bioscoop start over vijf minuten de film

Davids opa was regisseur, hij leerde hem veel over het maken van een film

Vanavond gaat ze naar de bioscoop, want er draait een leuke nieuwe film

Ze vond “Schindler’s List” een erg droevige film

Hij heeft veel horror gezien maar “The Ring” vond hij écht een enge film

Van de “Lord of the Rings”-reeks vond ze deel 1 de beste film

Het is zonde om naar de bioscoop te gaan om te kijken naar een geflopte film

Een acteur kan spelen in zowel een theater als een film

Op dit moment is Brad Pitt bezig met het maken van een nieuwe film

